# Fibronectin meshwork controls epithelial stem cell fate

**DOI:** 10.1101/2021.05.26.445735

**Authors:** Soline Estrach, Lionel Tosello, Floriane S. Tissot, Laurence Cailleteau, Ludovic Cervera, Kim B Jensen, Chloé C. Féral

## Abstract

Adult stem cell fate is tightly balanced by the local microenvironment called niche and sustain tissue regeneration^1–4^. How niche signals are integrated and regulate regeneration remains largely unexplored. The extracellular matrix and integrin ligand fibronectin is a crucial and well-characterized wound healing actor that has never been involved in skin regeneration. Here, we show that fibronectin displays a highly specific enrichment in hair follicle stem cells (HFSC) at the onset of regeneration. Conditional deletion of fibronectin in HFSC compartment (Lrig1, K19) leads to hair regeneration blockade, impaired stem cell location and fate. Dermal injection of exogenous fibronectin rescues these phenotypes. To elucidate molecular mechanism underlying fibronectin function, we used conditional deletion models of SLC3A2, the main integrin coreceptor. We show that, via its role in integrin-dependent assembly of fibronectin matrix, SLC3A2 acts as molecular relay of niche signals. Thus, fibronectin-integrin-SLC3A2 cascade finely tunes HFSC fate and tissue regenerative power.

## MAIN

Tissue homeostasis is achieved through a tightly controlled balance of growth and regression. To investigate physiological tissue regeneration, we use the mouse hair follicle^5^, which cycles between phases of growth and regression while maintaining a pool of stem cells (HFSC) to sustain tissue regeneration. HFSC via niche interactions fuel the hair regeneration cycle. Regulatory signals that balance SC quiescence and activity are provided by the niche. As a permanent part of the niche, extracellular matrix (ECM) components are key players of its instructive power in multiple organs^1–4^. In the skin, HFSC, residing in a specific location called the bulge^6^, are uniquely defined by expression of several markers such as integrin α6, CD34 but also keratin (K) K15, K19, LIM homeobox 2 (Lhx2), SOX9, leucine-rich repeat containing G protein-coupled receptor 5 (Lgr5)^7–12^ and leucine-rich repeats and immunoglobulin-like domains 1 (Lrig1). Even though, cell fate decisions are affected by signals from the ECM within the stem cell niche, how ECM composition and signalling are regulated remains unclear. Here, we unravel how specific niche signals are integrated in the bulge and how cell fate decisions are tuned by fibronectin signals.

To assess how ECM composition regulates regenerative signal in the niche, we analyzed specific enrichment of transcripts in stem cell compartment versus basal layer. We used Lrig1CreER^T2^GFP line to isolate GFP positive cells, that are located in the junctional zone in the upper isthmus that contribute to all 3 skin epithelial lineages in grafting experiments^13^. We isolated cells Lrig1 positive and integrin a6 positive (the basal population) using flow cytometry from day 28 (D28) back skin. D28 represents a specific time point when HFSC have received activation signal and start regenerative processes. Very interestingly, we observed a specific mRNA enrichment of fibronectin and Col17A1 (Fig. 1a). Col17A1 has previously been described^2,4^ as a crucial regulator of HFSC activity, validating our small-scale transcriptomic unbiased analysis. Next, we stained epidermal tail wholemount and very thick section of back skin for fibronectin expression. We observed a fibronectin meshwork encompassing the bulge area and the transit amplifying compartment as highlighted with brackets (Fig. 1b-e). Strikingly, no staining was observed in the interfollicular epidermis (IFE) leading us to speculate that the specific pattern observed is associated to a regenerative phenotype. We are convinced that this fibronectin signal was not detected so far in studies as only whole tissue imaging on tail or very thick 100 microns back skin sections allowed us to visualize it. We showed colocalization of the Lrig1^+^ cells with the fibronectin meshwork during the regenerative phase by epidermal wholemount staining (Fig. 1f). Using Imaris software, we reconstituted the 3D pattern of Lrig1 and fibronectin expression and revealed that Lrig1+ cells in regenerative phase rested on fibronectin tracks (Fig. 1g,h).

**Figure 1.**
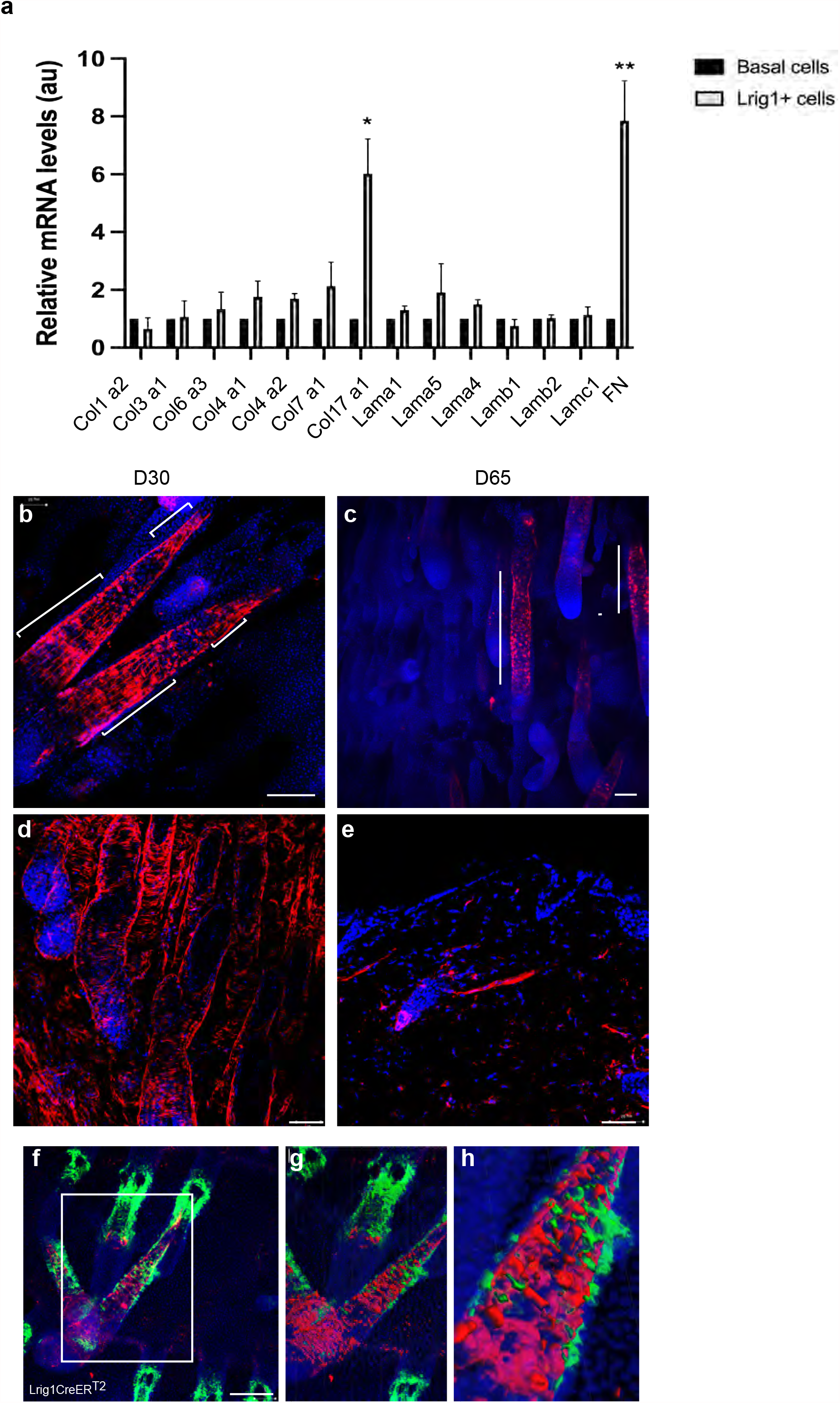
Fibronectin expression is enriched in the HFSC compartment during hair regeneration. a-Quantitative RT-PCR of ECM components from Lrig1+cells sorted and Basal cells from back skin at D28. b-c. Epidermal tail whole mount imaging with antibodies to fibronectin (red) from controls (vehicle-treated Lrig1CreER^T2^GFP, SLC3A2^fl/fl^) at D30 (b, confocal imaging scale bar 50 µm) and D65 (c, Light sheet imaging scale bar 50µm). d-e Immunostaining of thick back sections at D30 (d) or D65 (e) of controls with fibronectin antibody. Scale bar 50µm. f-h. Epidermal tail whole mount imaging with antibodies to fibronectin (red), GFP (green) from Lrig1CreER^T2^GFP, SLC3A2^fl/fl^ vehicle (Veh)-treated (f). Scale bar 50 µm; Imaris 3D volume rendering of the boxed area (g), enlarged area of M (h). Nucleus is stained with DAPI (blue).

To confirm that fibronectin is the ECM signal regulating regenerative potential of the HFSC, we generated conditional deletion of fibronectin (FN^fl/fl^) in epidermis using K19 and Lrig1 Cre lines, both inducible by 4OHT treatment. Lrig1CreERT2GFP is a GFP expression reporter line^13^. Fibronectin deletion was induced by 4OHT-topical application starting at D19 (Fig. 2a and Supplementary Fig. 1a) before endogenous signal of regeneration. We validated fibronectin deletion in both Lrig1CreER^T2^GFP and K19CreER lines by epidermal wholemount immunofluorescent staining (Fig. 2b,c; Supplementary Fig. 1b,c). Fibronectin deletion in both Lrig1 and K19 compartment led to a blockade in the first hair cycle during hair regeneration at D30 (Fig. 2d, Supplementary Fig. 1d), confirmed by H&E staining (Fig. 2e,f). Hair eventually regrew at D51 but, very interestingly, when hair was reclipped at D65 (resting phase of the hair cycle), before the start of the second synchronous hair cycle, the blockade reappeared and could not be bypassed anymore. One can speculate that some residual fibronectin was still present in the bulge inducing a delayed HFSC activation. We also observed an HF thinning at D158 in both Lrig1CreER^T2^GFP and K19CreER treated mice (Fig. 2g,h and Supplementary Fig. 1e,f) as indicated by arrow heads, strongly suggesting HFSC defects. Analysis of Lrig1 reporter expression in epidermal tail wholemount vehicle treated revealed expected staining in the junctional zone (adjacent to the sebaceous gland), bulge and bulbs. Interestingly, fibronectin deficient Lrig1^+^ cells were mainly located above the sebaceous gland, in a zone called infundibulum, starting at D30 (Fig. 2i,j), where they were still located at D158 (Fig. 2k,l). Moreover, the infundibulum displayed an enlarged morphology (2-fold larger than the controls) as visualized by Lightsheet microscopy imaging (Fig2. m-p, quantified in 2q). A similar enlargement was observed in the K19CreER deletion line (Supplementary Fig1. g,i). Flow cytometry analysis of the α6^+^ CD34^+^ population revealed that depletion of fibronectin driven by either Lrig1 or K19 promoter led to defects in the SC population and more specifically a significative reduction of the α6^High^ CD34^+^ population of SC (Fig. 2r, Supplementary Fig. 1j).

**Figure 2:**
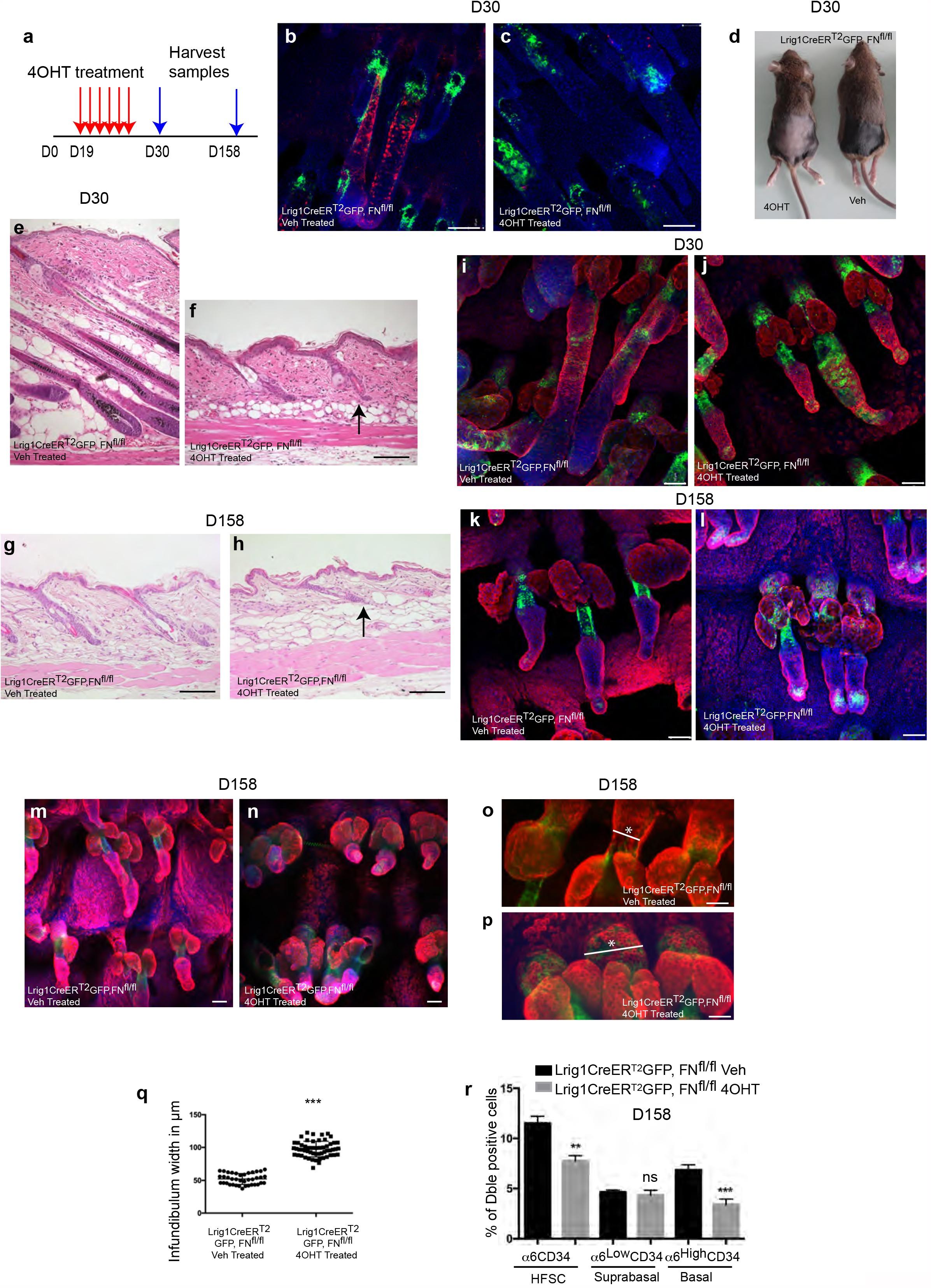
Fibronectin depletion in the stem cell compartment leads to hair regeneration blockade and HFSC deficient location. a Schematic representation of the experimental procedure used in panel b-r. b-c Epidermal tail whole mount imaging from vehicle-(Veh, b) or 4OHT-treated (c) of Lrig1CreER^T2^GFP, FN^fl/fl^ mice at D30 with antibody to FN (red) and GFP (green). Nuclei are in blue (DAPI). Scale bar 50µm. d-Picture of Lrig1CreER^T2^GFP, FN^fl/fl^ vehicle-(Veh) or 4OHT-treated at D30. Mice were clipped before the first topical treatment. e-h H&E staining of back skin section of Lrig1CreER^T2^GFP, FN^fl/fl^ mice at D30 (b, c) and D158 (d, e), Scale bar 100µm. i-l. Epidermal tail whole mount imaging with antibodies to GFP (green), K14 (red) from vehicle (veh)-or 4OHT-treated Lrig1CreER^T2^GFP, FN^fl/fl^ mice at D30 (i,j) and D158 (k, l). DAPI in blue. Scale bar 50µm. m-n Light sheet imaging of Lrig1CreER^T2^GFP, FN^fl/fl^ with indicated treatment. GFP (green), K14 (red), DAPI (blue). o-p Zoomed area showing infundibulum enlargement as indicated by the star. Scale bar 50µm. q quantification of infundibulum width in μm. (n=3; ****p≤ 0.0001). r. Quantification of indicated populations of epidermis from vehicle-(Veh) and 4OHT-treated Lrig1CreER^T2^GFP, FN^fl/fl^ mice labelled with antibodies to α6 integrin (α6) and CD34 and analysed by flow cytometry (n=6 per group; **p≤0.001; ***p≤ 0.0001).

To analyse more specifically HFSCs maintenance, we treated Lrig1CreER^T2^GFP mice at D65, corresponding to the resting phase of the hair cycle, then performed a chase for 7 months (D204) (Fig. 3a). We observed a total blockade of HF growth along with the thinning of the HF as indicated by arrow heads (Fig. 3b,c). Very interestingly, we saw the GFP+ cells defective location using both confocal (Fig3. d,e) and light sheet (Fig3. f,g) imaging and the infundibulum enlargement (Fig. 3h) as in mice treated at D19. Consistently, flow cytometry analysis revealed a depletion of the SC α6CD34 and more drastically of the α6^High^CD34^+^ compartment compared to controls (Fig. 3i). Altogether, those data suggest that fibronectin expression in the stem cell compartment is required for stem cell anchorage in the niche during hair regeneration.

**Figure 3:**
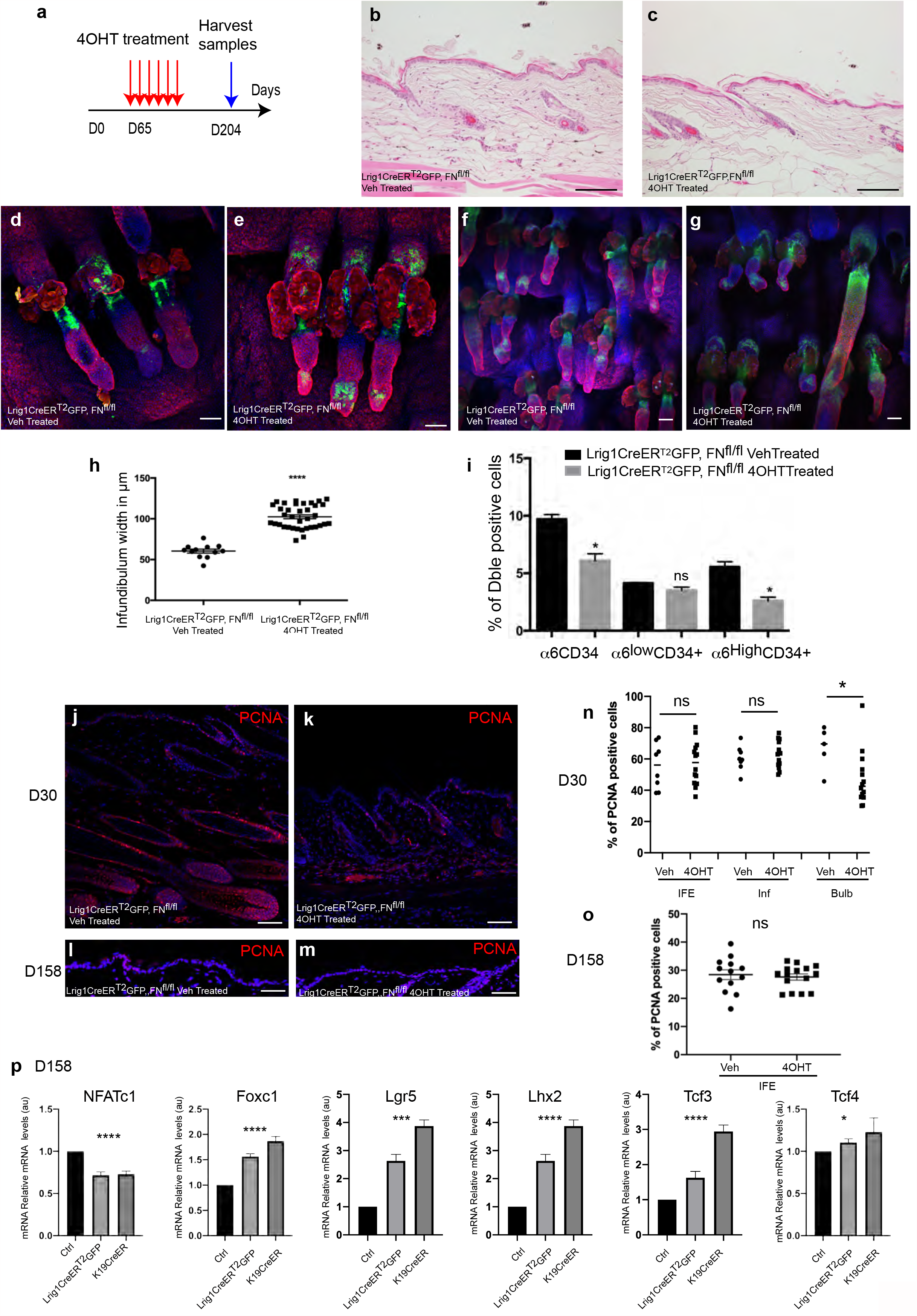
Fibronectin depletion in the stem cell compartment leads to defective HFSC signalling. a. Schematic representation of the experimental procedure used in panel b-i. b-c. H&E staining of back skin section of vehicle (Veh)-and 4OHT-treated Lrig1CreER^T2^GFP, FN^fl/fl^ mice at D204. Scale bar 100µm. d-g. Epidermal tail whole mount imaging with antibodies to GFP (green) and K14 (red) from vehicle (Veh)-and 4OHT-treated Lrig1CreER^T2^GFP, FN^fl/fl^ mice at D204. Scale bar 50µm. h. Quantification of infundibulum width in μm. (n=3). Nuclei in blue (DAPI) ****p≤ 0.0001. i-Quantification of indicated populations of epidermis from vehicle-(Veh) and 4OHT-treated Lrig1CreER^T2^GFP, FN^fl/fl^ mice labelled with antibodies to α6 integrin (α6) and CD34 and analysed by flow cytometry at D204 (n=6 per group). n.s. not significant. p values *p≤0.01). j-m. PCNA immunostaining of back skin sections at D30 (j, k) and D158 (l, m) of Lrig1CreER^T2^GFP, FN^fl/fl^ treated with vehicle (Veh) or 4OHT. Nuclei in blue (DAPI). Scale bar 50 µm. n, o Percentage of PCNA positive cells in the different compartment of the epidermis at D30 (n) and at D158 (o). IFE interfollicular epidermis, Inf infundibulum, Bulb. p-Quantitative RT-PCR of Stem Cell markers (Lef1, Nfatc1, Foxc1, Lgr5, Lhx2, Tcf3 and Tcf4) from back skin RNA freshly isolated nd D158 (H). Data shown are mean ±SEM (n=6 per group for Lrig1CreER^T2^GFP, FN^fl/fl^ and K19 CreER, FN^fl/fl^ vehicle-(Veh) or 4OHT-treated, (*p≤0.01, ***p≤ 0.0001, ****p≤ 0.00001, n.s. not significant).

To decipher the molecular signals impaired upon fibronectin deletion in K19 and Lrig1 compartment, we analysed the proliferation status of the epidermis at both D30 (Fig. 3j,k,n) and D158 (Fig. 3l,m,o). Bulb proliferation was strongly affected at D30 and can be directly correlated with the hair growth blockade phenotype. There was no other defect in cell proliferation in Lrig1CreER^T2^GFP epidermal compartment following fibronectin deletion at D30 and D158 (Fig. 3n,o).

We then went on looking for the signalling pathways involved in the hair regeneration cycle such as Wnt and BMP^14^. At D30, Lef1 activation was defective as expected since no HF growth was observed. RTqPCR analysis of BMP ligands and BMP target genes (Id) revealed no defects in this pathway, involved in SC quiescence maintenance (Supplementary Fig. 2a). Interestingly, HFSC deficient in fibronectin at D30 displayed a strong induction of NFATc1 reflecting initiation of HFSC activation process^15^, nonetheless this activation was incomplete and didn’t lead to HF growth (Fig. 2d; Supplementary Fig. 2a). Lrig1CreER^T2^GFP, FN^fl/fl^ 4OHT-treated mice showed increased stem cell markers at D158 (Fig. 3p). Strikingly, they also presented high level of Foxc1 mRNA^16,17^ indicating that the increased pool of HFSC remained in a quiescent state^17^. BMP signalling was also not affected at D158 either (Supplementary Fig. 2b). K19CreER, FN^fl/fl^ mice at D158 presented the same results on stem cell markers (Fig. 3p, Supplementary Fig. 2b). Thus, fibronectin deletion in HFSC could result into an accumulation of cells retaining specific HFSC markers outside of the niche as the matrix specific anchoring signal is deficient. Flow cytometry and RTqPCR data seemed inconsistent as a6 levels are reduced but it has been described that integrin levels are controlled by ECM ligands expression^18^. Hence, one can speculate that in absence of fibronectin expression a6 expression decreased. In conclusion, fibronectin contributes to complete activation of stem cell in order to fuel HF regeneration.

If fibronectin is sufficient to fuel hair regeneration, adding exogenous fibronectin into the dermis of HFSC deficient mice should rescue hair growth. Lrig1CreER^T2^GFP, FN^fl/fl^ mice were treated with 4OHT at D19 then dermally injected either with PBS or soluble fibronectin at D24. We followed the first synchronous hair cycle and monitored hair growth (Fig. 4a-o). Skin gets darker when HF growth is effective (Fig. 4a). When comparing 4OHT-treated Lrig1CreER^T2^GFP, FN^fl/fl^, we observed darker spots only when exogenous fibronectin was injected compared to PBS-injected back-skin which remained pink (Fig. 4c vs 4b). PBS had no positive effect on HF regeneration. Exclusively exogenous fibronectin injection (compared to PBS) rescued hair growth in Lrig1CreER^T2^GFP, FN^fl/fl^ 4OHT-treated mice (Fig. 4d,f). Moreover, it restored proliferation in the bulb as well as GFP expression pattern in regenerating HF (Fig. 4g-l). We confirmed, using thick back skin sections, that exogenous fibronectin was assembled in 4OHT-treated Lrig1CreER^T2^GFP, FN^fl/fl^ mice (Fig. 4m-o). In conclusion, exogenous injection of fibronectin rescues HF regeneration in mice deficient for fibronectin in HFSC.

**Figure 4:**
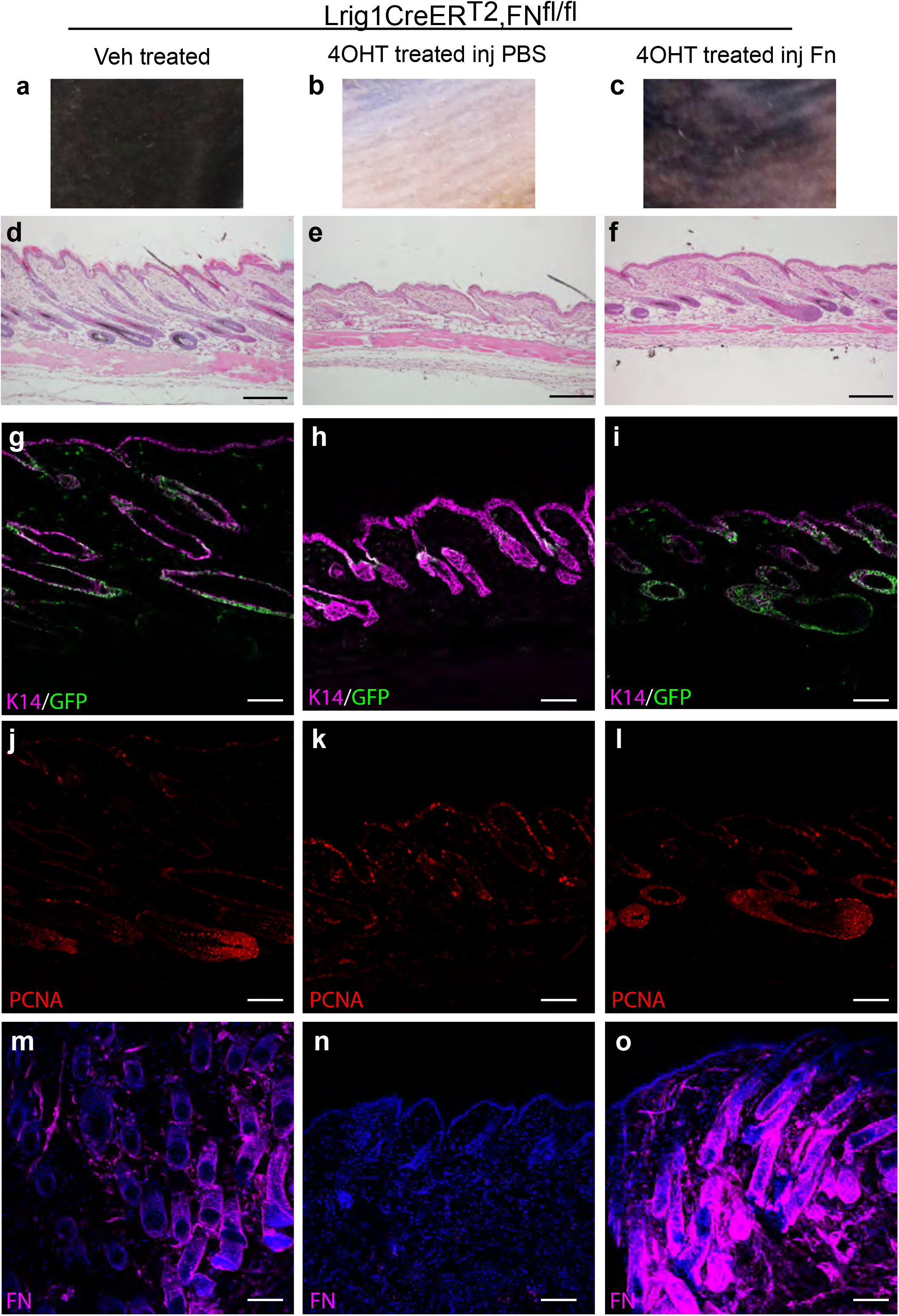
Fibronectin signalling is required and sufficient for hair follicle regeneration. a-c Pictures of back skin collected from Lrig1CreER^T2^GFP, FN^fl/fl^, vehicle (Veh) treated (a), 4OHT-treated then injected with either PBS (b) or 60 μg of FN (c). d-f. H&E staining of back skin sections of mice indicated above. Scale bar 100µm. g-o. Immunostaining of 5μm-thick back skin sections with antibody to K14(magenta)/GFP (green) (g, h, i) and PCNA (red) (j, k, l). m-o-Immunostaining of corresponding 100μm-thick back skin sections with antibody to FN. Nuclei in blue (DAPI). Scale bar 50µm. p. Schematic representation of the epidermal reconstitution assay performed. q. Representative grafted area. r. Grafted area were imaged from the dermal side using a stereomicroscope coupled to fluorescence imaging. s. H&E staining of representative sections of grafted area. Scale bar 100 µm

Interestingly, in the skin, fibronectin has only been described and observed during wound closure. A granulation tissue is generated in order to form a provisional matrix that will enable keratinocytes migration and repair^19^. Fibronectin is one main component of this structure which disappears once the reepithelialisation has occurred. The neodermis is formed with no fibronectin expressed leading to HFSC reprogramming into interfollicular fate^20^. Here, we clearly demonstrate that fibronectin controls adult stem cells in the skin.

Next, we ought to get a better insight into the signalling mechanisms involved. Fibronectin is a ligand for ECM receptors, integrins. Integrin-dependent signals play a crucial role in stem cell maintenance. ECM ligand binding is required for maintenance of regenerative capacity of adult stem cells^2–4^. Thus, we hypothesized that SLC3A2, the main integrin co-receptor via its capacity to enhance integrin signalling and to control fibronectin assembly^21^, acts as the molecular relay of the niche signal. To assess this hypothesis, we first evaluated SLC3A2 level of expression in bulge stem cells using MFI from flow cytometry data of α6/CD34 populations (Fig. 5a) obtained from WT resting back skin. SLC3A2 is strongly expressed in both basal and suprabasal HFSC (Fig. 5b). As expected, α6 positive basal keratinocytes also express SLC3A2 level^22^. Second, we decided to especially knock-out *SLC3A2* in the HFSC using the same K19 and Lrig1 Cre lines and treatment as previously (Fig. 5c). Recombination efficacy was confirmed by flow cytometry (Lrig1, K19) (Supplementary Fig. 3a,b;d,e).

**Figure 5.**
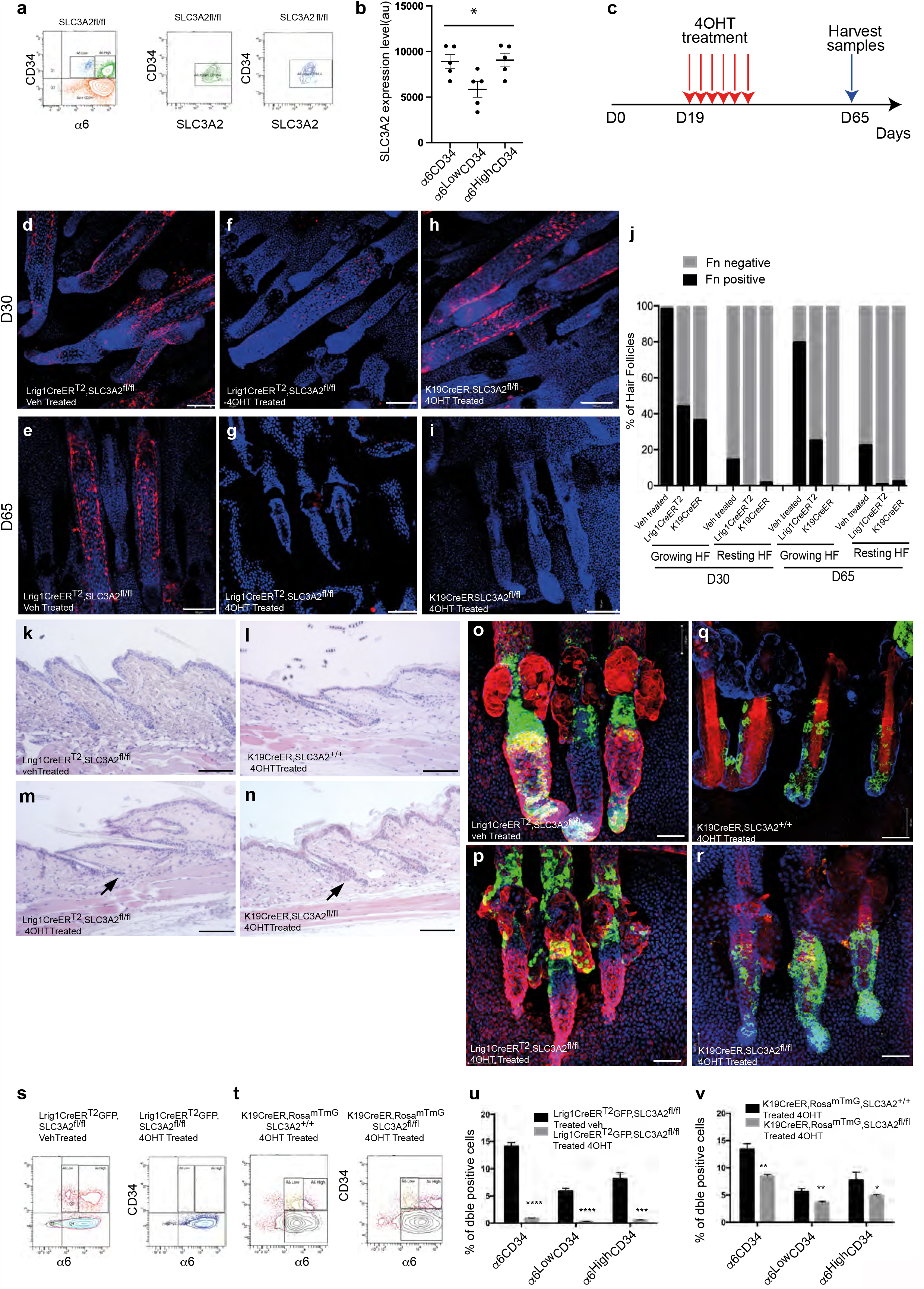
SLC3A2 depletion in the epidermal stem cell compartment results in defective epidermal stem cells fate. a-Flow cytometry analysis of SLC3A2 expression in fresly isolated keratinocytes depicted in colour, HFSC (α6^High^CD34^+^, green and α6^low^CD34^+^, blue). Plots representing SLC3A2 level of expression are colour coded. b Quantification of fluorescence intensity using Geomean data from panel a. (HFSC hair follicle stem cells, ANOVA ***p≤0.001). c. Schematic representation of experimental procedure used in panel d-v. d-i. Epidermal tail whole mount imaging at D30 or D65 with antibody to fibronectin (red) either vehicle-(veh, d, e) or 4OHT-(f, g, h, i) treated mice (lines as indicated). Scale bar 100 µm. j. Quantification of FN positive staining on hair follicle elongated according to the phase of the regenerative cycle. k-n. H&E staining of representative D65 back sections of vehicle(veh)-(k) or 4OHT-treated Lrig1CreER^T2^GFP, SLC3A2^fl/fl^(m); K19CreER, Rosa^mTmG^, SLC3A2^+/+^(l), K19CreER, Rosa^mTmG^, SLC3A2^fl/fl^ (n) mice. Scale Bar 100 μm. o-r. D65 Epidermal tail whole mount imaging with antibodies to GFP (green) and K14 (red) from Lrig1CreER^T2^GFP, SLC3A2^fl/fl^ (o, p); K19CreER, Rosa^mTmG^, SLC3A2^+/+^(q), K19CreER, Rosa^mTmG^, SLC3A2^fl/fl^ (r) mice with indicated treatment (veh, vehicle or 4OHT). Nuclei staining in blue (DAPI). Scale bar 50 µm. s-v. Flow cytometry analysis in telogen epidermis of α6^+^CD34^+^ populations, in vehicle-(veh) or 4OHT-treated Lrig1CreER^T2^GFP, SLC3A2^fl/fl^ (s); and 4OHT-treated K19CreER, Rosa^mTmG^, SLC3A2^+/+^, K19CreER, Rosa^mTmG^, SLC3A2^fl/fl^ (t). u-v. Quantification of the population percentage from flow cytometry analysis for Lrig1 and K19 lines respectively (n=3 per group; ****p≤0.0001; ***p≤0.001; **p≤;0.01;*p≤0.05).

We then analysed fibronectin pattern of expression in the SLC3A2 stem cell deficient models to correlate defective integrin signalling, fibronectin meshwork and HF regeneration (Fig. 5d-i). We scored fibronectin staining present on HF at D30 and D65 (Fig. 5d-e) in both SLC3A2 stem cell KO models (Lrig1 Fig. 5f,g and K19 Fig. 5h,i) on epidermal wholemount. While growing HF of vehicle-treated mice presented a strong fibronectin meshwork signal, the very limited number of growing HF of 4OHT-treated mice displayed a drastic reduction of fibronectin expression (Fig. 5j). Defects in fibronectin staining observed on tail wholemount were also observed on back skin sections (data not shown).

Deletion of SLC3A2 under Lrig1 and K19 promoter resulted in major blockade of HF regenerative cycle (Supplementary Fig. 3c,f) compared to vehicle-treated mice, confirmed by H&E staining (data not shown). Phenotypes observed in mice deficient for SLC3A2 in Lrig1 and K19 compartments were drastic with only 16% and 26% HF growth, respectively, compared to vehicle-treated mice as quantified on back skin sections and epidermal tail wholemount (Supplementary Fig. 3g). The same scoring was performed on epidermal tail whole mount D65 samples where only a limited number of HF was still elongated in control mice (22%) and almost none in stem cell compartment knock out mice (Supplementary Fig. 3h). This HF phenotype worsened with time as observed at the first synchronous resting phase (D65) using H&E staining (Fig. 5k-n). The bulbs displayed a very thinned and abnormal morphology in Lrig1CreERT^2^GFP (Fig. 5k,m) as indicated by arrow heads. In the K19CreER, Rosa^mTmG^ line, the bulge exhibited an enlarged morphology with abnormal bulbs (Fig. 5l,n). Next, we used GFP and Rosa^mTmG^ recombination for tracing cells in the Lrig1 and K19 lines respectively. Defective location of GFP positive cells was extremely severe in Lrig1-and K19-driven deletion of SLC3A2 (Fig. 5o,r). SLC3A2-deficient-Lrig1+ cells displayed mainly a prominent location in the infundibulum and depletion from the bulbs (Fig. 5o vs 5p). Lineage tracing experiments for K19CreER, Rosa^mTmG^ control mice revealed expected staining in the bulge and bulbs at D65 on epidermal wholemount (Fig. 5q). Interestingly, in K19CreER, Rosa^mTmG^, SLC3A2^fl/fl^ 4OHT treated mice, we observed strong labelling in the bulge and in the bulb that didn’t result into hair growth (Fig. 5q vs 5r). Flow cytometry analysis of SC in both Lrig1CreERT^2^GFP and K19CreER, Rosa^mTmG^ mice at D65 revealed a drastic loss of the α6^+^ CD34^+^ double positive population (Fig. 5s and 5t respectively, quantified in 5u and 5v). SLC3A2 deletion in HFSC displayed the same HFSC defects but a stronger phenotype than the fibronectin one. This could be directly linked to the role of downstream molecular relay played by SLC3A2. SLC3A2 is downstream of the ECM ligand, modulates the signal from the integrin receptors that mediate several signals^23^. Those signals regulate cell shape, cell proliferation, actin cytoskeleton thus amplifying the unique initial signal from the specific ECM ligand fibronectin to several target signalling.

RT-qPCR analysis for SC markers Lhx2, Lgr5 and Tcf3, confirmed flow cytometry results (Supplementary Fig. 3i) and suggest that HFSC rely on SLC3A2 for their fate.

To verify this hypothesis, we performed grafting experiments. We used freshly isolated FACS-sorted keratinocyte subsets from Lrig1CreERT^2^GFP, SLC3A2^fl/fl^ mice to test the ability of this subpopulation of HFSC to regenerate HF in vivo. We injected GFP+ cells from untreated mice in grafting chamber (in combination with P2 dermal fibroblasts) on the back of *Nude* mice. One week later, to avoid interfering with grafting, we topically treated the graft with 4OHT or vehicle (Supplementary Fig. 3j). Then, we monitored HF growth over a 6-week period, at which time, we harvested the samples. Our data show that, while vehicle treated Lrig1CreER^T2^GFP, SLC3A2^fl/fl^ cells sustained HF regeneration, SLC3A2 deficiency totally blocked SC potential to reconstitute HF (Supplementary Fig. 3k). Only vehicle-treated grafts showed GFP bulbs (Supplementary Fig. 3l). No HF grew out of 4OHT-treated grafts (Supplementary Fig3.l). H&E staining confirmed the absence of HF growth in grafted area of 4OHT-treated mice compared to vehicle-treated (Supplementary Fig. 3m). These results show that SLC3A2 is necessary for HFSC cell autonomous function *in vivo*.

In conclusion, deletion of SLC3A2 in HFSC compartment disrupt fibronectin meshwork assembly during hair follicle regeneration. Thus fibronectin-integrin-SLC3A2 signalling cascade is necessary and required for HFSC regenerative capacity.

Altogether, our data highlight the unique role of a specific ECM component of the niche, namely fibronectin, in HFSC activation. In accordance with recent data on maintenance of ES cells pluripotency by fibronectin^24^, our results demonstrate that it plays a crucial role in adult stem cells maintenance too. On a more conceptual point of view, we show that ECM specific anchoring signal via fibronectin tunes HFSC fate and regenerative power of this population. Research on epithelial regeneration, specifically in the skin, will now be shaped by this novel finding of a niche-induced fibronectin meshwork controlling SC renewal and tissue regeneration.

## Acknowledgments

The authors thank Dr E.Boulter and Dr L. Seguin for careful reading and meaningful insights. The authors acknowledge the IRCAN CytoMed, PICMI, GenoMed and Animal core facilities which are supported by grants from the Conseil Général 06, the FEDER, the Ministère de l’Enseignement Supérieur, the Région Provence Alpes-Côte d’Azur, the Canceropole PACA, the foundation ARC and INSERM. This study was supported by grants from the InCa (R19004AP), la Fondation ARC (R18020AA) and. FST was the recipient of a doctoral fellowship from INSERM Region Provence-Alpes Cote d’Azur Canceropole PACA. This work was supported by the French Government (National Research Agency, ANR) through the “Investments for the Future” LABEX SIGNALIFE: program reference UCAJEDI ref. # ANR-15-IDEX-01.

