## Supplemental Data for "Fibronectin meshwork controls epithelial stem cell fate" (from another version of this preprint: version 1, 2021-05-26)

**Supplementary Figure legends**

*Supplementary Figure 1: Fibronectin depletion in K19+ compartment leads to hair regeneration blockade and HFSC deficient location.*

a. Schematic representation of the experimental procedure used in panel b-j. b-c Epidermal tail whole mount imaging from vehicle-(Veh, b) or 4OHT-treated (c) of K19CreER, $FN^{fl/fl}$ mice at D158 with antibody to FN (red) and GFP (green). Nuclei are in blue (DAPI). Scale bar 50$\mu$m. d-Picture of K19CreER, $FN^{fl/fl}$ vehicle-(Veh) or 4OHT-treated at D30. Mice were clipped before the first topical treatment. e-f H&E staining of back skin section of K19CreER, $FN^{fl/fl}$ mice at D158, Scale bar 100$\mu$m. g-h. Epidermal tail whole mount imaging with antibodies to K14 (red) from vehicle (veh)- or 4OHT-treated K19CreER, $FN^{fl/fl}$ mice at D158 (g,h). DAPI in blue. Scale bar 50$\mu$m. Infundibulum is indicated by stars. i Quantification of infundibulum width in µm. (n=3; *p≤ 0.05). j. Quantification of indicated populations of epidermis from vehicle- (Veh) and 4OHT-treated K19CreER, $FN^{fl/fl}$ mice labelled with antibodies to α6 integrin (α6) and CD34 and analysed by flow cytometry (n=3 per group; **p≤0.001).

*Supplementary Fig 2: Molecular Pathway analysis of D30 and D158 epidermis*

a- Quantitative RT-PCR of Stem Cell markers (Bmp4, Bmp6, Foxc1, Lef1, Nfatc1, Id1,Id2,Id3) from back skin RNA freshly isolated from Lrig1CreER$^{T2}$GFP, $FN^{fl/fl}$ at D30 (A)

b- Quantitative RT-PCR of Stem Cell markers (Bmp4, Bmp6, Id1,Id2,Id3) from back skin RNA freshly isolated from Lrig1CreER$^{T2}$GFP, $FN^{fl/fl}$ at D158 and (Bmp4, Bmp6) from K19CreER$^{T2}$, $FN^{fl/fl}$ at D158

Data shown are mean ±SEM (n=4 per group at D30 and n=6 at D158). Data shown are mean ±SEM (n=4 per group). n.s. not significant, *p≤0.01, ***p≤ 0.0001.

*Supplementary Fig 3: SLC3A2 deletion in HFSC compartment impairs SC function.*

. a. Analysis of deletion efficacy in Lrig1 CreER$^{T2}$GFP using Flow cytometry. b Quantification of deletion efficacy in Lrig1 CreER$^{T2}$GFP. c. Pictures of Lrig1Cre ER$^{T2}$GFP, SLC3A2$^{fl/fl}$ vehicle or 4OHT treated as indicated at D30. d. Analysis of deletion efficacy in K19 CreER using Flow cytometry. e Quantification of deletion efficacy in K19 CreER. f. Pictures of K19Cre ER, SLC3A2$^{+/+}$; K19Cre ER, SLC3A2$^{fl/fl}$ 4OHT treated as indicated at D30. g- Quantification of the percentage of hair follicle (HF) on back skin in growth phase (D30) 200 HF were scored per mouse line and one-way ANOVA statistical analysis performed ****p≤ 0.0001. h- Quantification of the percentage of hair follicle (HF) in growth phase (D30) and in resting phase (D65) on epidermal wholemount. 200 HF were scored per mouse line.

i- Quantitative RT-PCR of Stem Cell markers (Tcf3, Lhx2, Lgr5, Sox9) from back skin of Lrig1CreER$^{T2}$GFP, SLC3A2$^{fl/fl}$ vehicle (Ctrl) or 4OHT treated; K19CreER, Rosa$^{mTmG}$, SLC3A2$^{+/+}$, K19CreER, Rosa$^{mTmG}$, SLC3A2$^{fl/fl}$ 4OHT treated mice. Data shown are mean ± SEM (n=15 for Lrig1CreER$^{T2}$GFP and K19CreER, Rosa$^{mTmG}$). HFSC *p≤ 0.01; **p≤ 0.001, ***p≤ 0.0001; ****p≤ 0.00001). j. Schematic representation of the epidermal reconstitution assay performed using Lrig1CreERGFP, SLC3A2$^{fl/fl}$ cells. k. Representative grafted area. l. Grafted area were imaged from the dermal side using a stereomicroscope coupled to fluorescence imaging. m. H&E staining of representative sections of grafted area. Scale bar 100 $\mu$m

.

**Experimental Procedures:**

**Mice**

All procedures CIEPAL#191 and CIEPAL#459 were approved by the Institutional Animal Care and Use Committee at the University of Nice-Sophia Antipolis, Nice, FR. Fibronectin conditional null mice, $Fn^{fl/fl}$ (Jackson Laboratory #029624) were crossed with Lrig1-EGFP-ires-$CreER^{T2}$ and K19 *CreER*. SLC3A2 conditional null mice, $SLC3A2^{fl/fl}$ [1] were crossed with Lrig1-EGFP-ires-$CreER^{T2}$ refer to as Lrig1CreER$^{T2}$GFP(kind gift from K.B Jensen, BRIC, Copenhagen) [2], K19 *CreER* (kind gift from G.Gu, Vanderbilt, USA) [3].

Littermates were assigned to experimental groups regardless of sex. No randomization of animals was performed. All quantifications were performed blinded as the identity of the sample was concealed from the staff performing the experiment. Monte-Carlo simulation was used to determine group size.

4OHT tamoxifen (#H7904 Sigma) was resuspended in Ethanol 100% at a concentration of 7,5mg/ml. Topical treatment (back and/or Tail) were performed as indicated (6 treatments of 1.5 mg 4OHT in ethanol), Ctrl groups were treated with vehicle (Ethanol).

**Histology and Immunohistochemistry.**

Tissue samples were fixed o/n in 3.7% formaldehyde, then washed in EtOH70% for 24h. Samples were then processed and embedded in paraffin. 7µm sections were produced. Formalin-fixed paraffin sections were used for all histology.

Hematoxylin and Eosin staining was performed as follow. Paraffin was removed using xylene 2x5 min, then sections were rehydrated via immersion 5 min for each of the following ethanol solution: 100%, 95%, 75%, 50% then dipped in distilled water for 5min; stained for 5 min in hematoxylin (S3309, DAKO), rinsed with tap water, 10 sec in NH4OH 0.08%, rinsed with tap water, stained for 30 sec in eosin (#HT1101166-500ml, Sigma-Aldrich), rinsed with tap water. Sections were dehydrated via immersion 2 min for each of the following ethanol solution 50%, 75%, 95%, 100%, 2x 2 min in xylene, Mount using non-aqueous mounting medium.

All immunostainings on sections were performed as follow, antigen retrieval was performed using citrate buffer 0.1M pH6, then sections were blocked in 5% Goat serum and 1%BSA, incubated with primary antibody in blocking solution (chicken anti-GFP (Aves lab Inc, USA)1:3000) o/n. Sections were washed 3 times 10 min in PBSTriton 0.01% then incubated with secondary antibody anti chicken alexa 488 (A-11039, Thermoscientific) in blocking solution then washed 3 times in PBS/Tween and once in water before mounting using Mowiol.

Mouse epidermal wholemounts were prepared and processed as previously described (Braun et al, 2003). Deboned tails were incubated in PBS 5mM EDTA for 4h at 37°C. Then epidermis is peeled of and fixed in 4%PFA at room temperature for 1h30 min. Epidermal samples were saturated in 0.9%NaCl, 20mM HEPES,0.5% skim milk powder, 0.5% TritonX-100 for 30 min at room temperature. Primary antibodies (rabbit anti-mouse cytokeratin 14 (PRB-155P, Covance):1:1000, chicken anti-GFP (Aves lab Inc, USA)1:3000, Fibronectin (Sigma)1:100) were diluted in blocking solution and incubated o/n at room temperature on a rocking table. Samples were washed 3 or 4 times at room temperature for 5h in PBS,0.2%Tween. Secondary antibodies (Alexa Fluor 488 or 594 or 647–conjugated anti–rabbit, chicken (Thermo):1:400) and DAPI were diluted in blocking solution and incubated o/n at room temperature on a rocking table. Samples were washed 3 or 4 times at room temperature for 5h in PBS,0.2%Tween. Then they were mounted using Mowiol. Imaging was performed using a Zeiss LSM 5 EXCITER confocal microscope.

**qPCR/RNA preparation:**

Total RNAs were extracted from back skin samples with Trizol reagent (GIBCO) using GentleMACS M tubes and the GentleMacs Dissociator (Miltenyi). Reverse transcription was performed on the same amount of total RNA with Superscript IV reverse transcriptase (Life, 18090050), using OligodT according to manufacturer's instructions. Real time PCR was performed using Fast SYBRgreen master mix (AppliedBiosystems) on a StepOne RealtimePCR thermocycler (ThermoFisher scientific). Primers for specific genes are listed in supplemental table1. Results were normalized to rplp0 using the ΔCt method. Statistical significance was determined with Student's t-test.

**Flow cytometry.**

Mouse Back skin was dissected and incubated by floating on trypsin 2.5% for 1h30 min at 37°C. Epidermis was separated from dermis then minced using scalpels. FAD low$Ca^{2+}$ **(**FAD low$Ca^{2+}$mixture of 3 parts $Ca^{2+}$-free DMEM and 1-part $Ca^{2+}$-free Ham's F12 supplemented with 1.8~$10^{-4}$ M adenine, hydrocortisone 0.4 µg/ml, cholera toxin $10^{-10}$ M, EGF 10 ng/ml, insulin 5µg/ml, Triiodothyronine 2.$10^{-9}$M, 10% FCS $Ca^{2+}$-free**)** was added to the minced epidermis. Then, pipet up and down the suspension until small clumps disaggregate. Cell suspension was filtered on cell strainer 70um then spin down 1500 rpm for 5 min. The cellular pellet was resuspended in PBS /2% FCS $Ca^{2+}$-free. Single cell suspension from GFP lines were directly sorted. Single cell suspension from adult epidermis were immunolabeled using antibodies to α6 integrin-PECy7 (1:500; clone G0H3, ebioscience), anti-CD34-BV421 (1:100; cloneRAM34, BD) and anti-CD98hc (1:100; clone RL388, BD). Samples were sorted or analyzed on a FACSARIAIII; with DiVA software (BD).

**Fibronectin injection:**

Mice Lrig1-EGFP-ires-*CreER$^{T2}$, FN$^{fl/fl}$ were clipped then treated 6 times with 4OHT from D19 on the back. At D26, 4 intradermal injection of 15µg each of fibronectin (Sigma) were performed using 29G syringe on the back. Hair growth was monitored by checking skin pigmentation. Mice were sacrificed at D34 when dark spots appeared on back skin. Then, samples were harvested.*

**Hair reconstitution Assay.**

Assay was performed as described [4]. Balb/c$^{nu/nu}$ mouse were anaesthetized using isoflurane, a 6mm punch was performed on the back where a 6mm silicon chamber was inserted. The chamber was secure using two wound staples.

Keratinocytes were isolated as described above. Dermal fibroblasts were prepared from P2 Balb/c pups. Skin was dissected and incubated by floating on Trypsin/Dispase solution for 1h at 37°C. After 1h, the epidermis was discarded and the dermis minced in small pieces in a 0,25% collagenase/ FAD lowCa$^{2+}$ solution for 1h at 37°C. Vigorously pipet the solution up and down then filtered on cell strainer, spinned down 5 min at 1500rpm then the pellet was washed once with FAD lowCa$^{2+}$ and resuspended in FAD lowCa$^{2+}$ for grafting.

Lrig1+cells were isolated from 8w-old mouse back skin and sorted by Flow cytometry using GFP fluorescence. $5x10^6$ dermal cells, $10^5$ Lrig1-GFP+ sorted epidermal keratinocytes and $3x10^6$ unfractionated, GFP-negative epidermal keratinocytes were implanted in the grafting chamber on the back of a Balb/c nude mouse. One week after grafting, three 4OHT treatment were performed (1.5mg in ethanol) in the grafted-chamber. Samples were harvested 5 weeks after.

**Statistical analysis**

For animal experiments, Monte-Carlo simulation was used to determine group size.

All animal experiments, Flow cytometry and RT-qPCR were performed in a blinded fashion. All quantifications represent mean ± standard error of the mean (s.e.m.). Images are representative of experiments that have been repeated at least three times. Group comparison was performed using two-tailed unpaired Student's t test. Multiple groups comparison was performed by 2way-ANOVA.

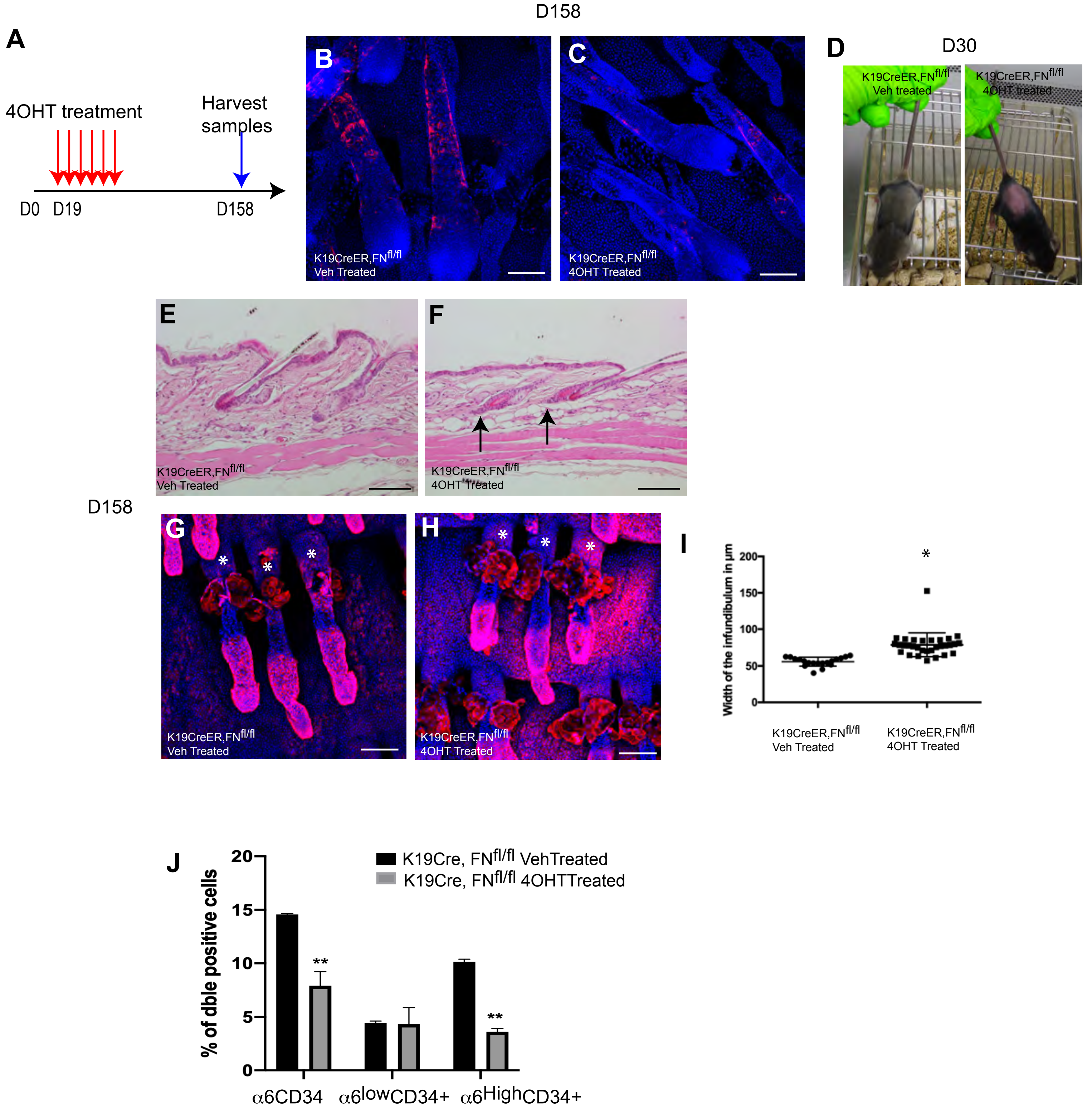
D158
A
4OHT treatment
Harvest samples
D0
D19
D158
B
K19CreER,FNfl/fl
Veh Treated
C
K19CreER,FNfl/fl
4OHT Treated
D
D30
K19CreER,FNfl/fl
Veh treated
K19CreER,FNfl/fl
4OHT treated
E
K19CreER,FNfl/fl
Veh Treated
F
K19CreER,FNfl/fl
4OHT Treated
D158
G
K19CreER,FNfl/fl
Veh Treated
H
K19CreER,FNfl/fl
4OHT Treated
I
Width of the infundibulum in µm
K19CreER,FNfl/fl
Veh Treated
K19CreER,FNfl/fl
4OHT Treated
J
% of dble positive cells
K19Cre, FNfl/fl VehTreated
K19Cre, FNfl/fl 4OHTTreated
α6CD34
α6lowCD34+
α6HighCD34+

Supp Fig1

**a**

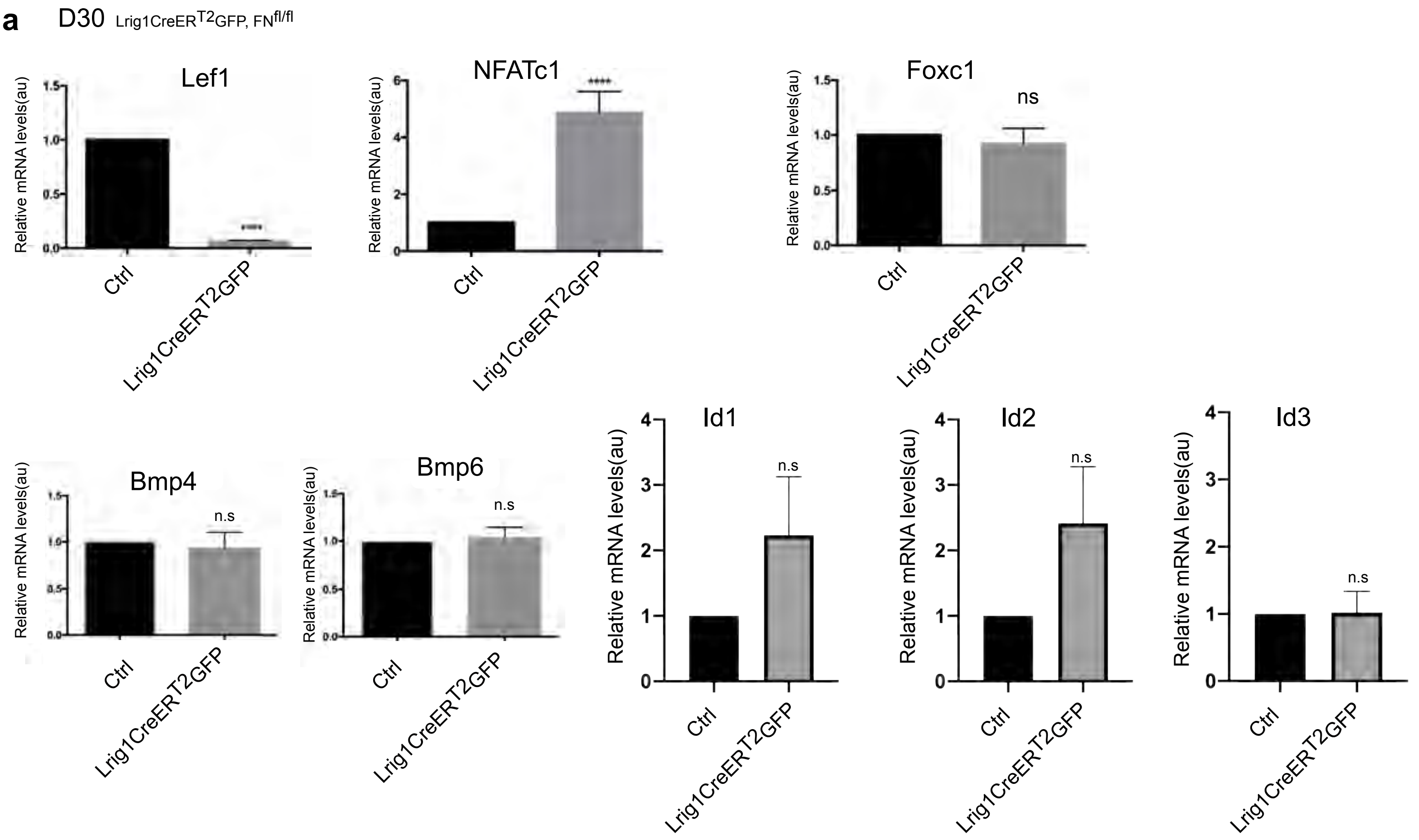


**b**

D158

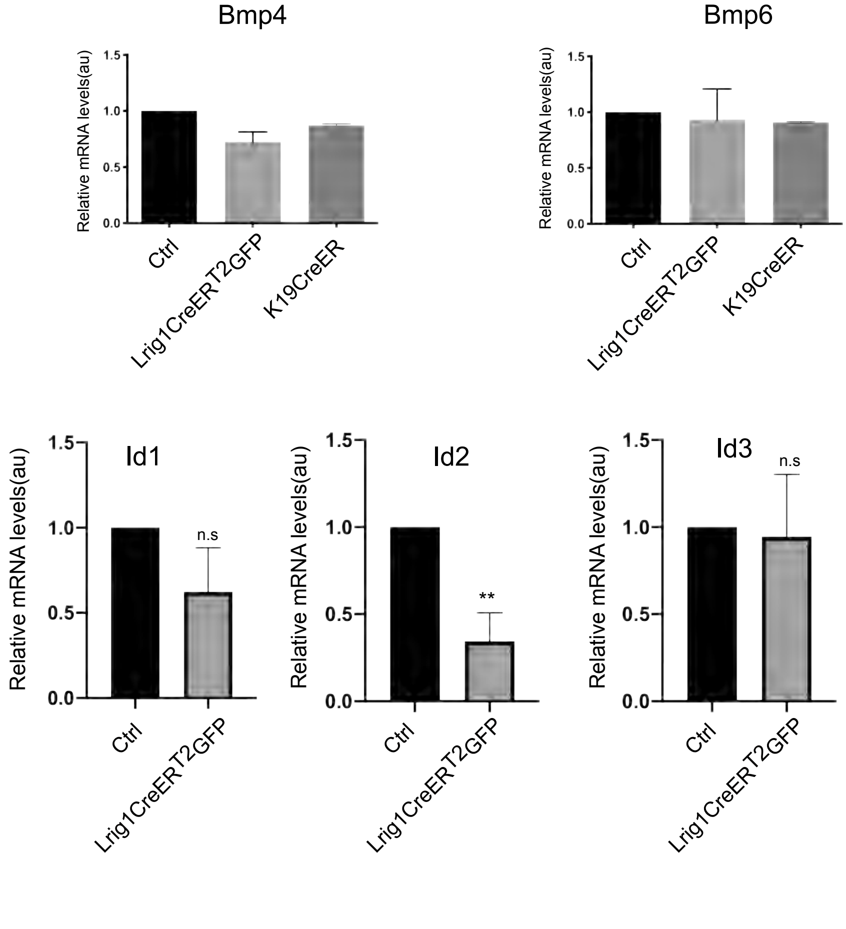


Supplementary Fig 2

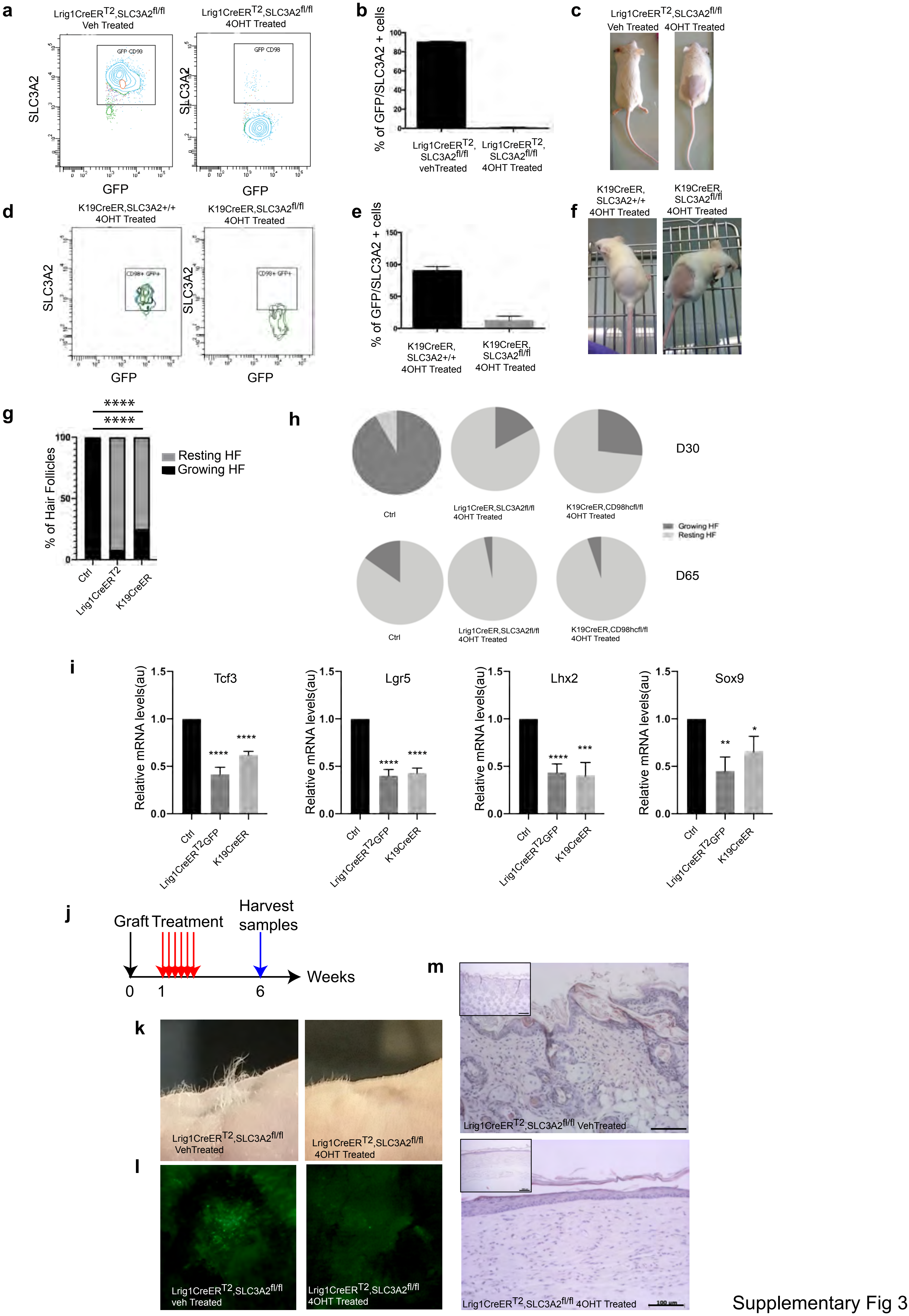


Supplementary Fig 3
